# A Scalable Framework for Identifying Allelic Series from Summary Statistics

**DOI:** 10.1101/2024.10.31.621375

**Authors:** Zachary R McCaw, Jianhui Gao, Rounak Dey, Simon Tucker, Yiyan Zhang, insitro Research Team, Jessica Gronsbell, Xihao Li, Emily Fox, Colm O’Dushlaine, Thomas W Soare

**Affiliations:** insitro, South San Francisco, CA, USA; Department of Statistics, University of Toronto, Ontario, CA; Department of Biostatistics, University of North Carolina at Chapel Hill, NC, USA; Department of Genetics, University of North Carolina at Chapel Hill, NC, USA

**Keywords:** Allelic series, Rare variant association testing, Variant Pathogenicity, Whole exome sequencing

## Abstract

Genes for which a dose-response relationship exists between mutational severity and phenotypic impact make for logical therapeutic targets, as the effects of pharmacological modulation can be anticipated from the natural variation present in a population. We refer to genes where such a dose-response relationship exists as harboring allelic series, and have introduced the rare coding-variant allelic series test (COAST) for their identification. The original COAST required access to individual-level data. However, such data are often unavailable due to privacy concerns or logistical challenges. Meanwhile, single-variant summary statistics of the type produced by genome-wide association studies are plentiful. Here we introduce COAST-SS, an extension of COAST that accepts summary statistics as input, namely the per-variant effect sizes and standard errors, along with estimates of the minor allele frequency and local linkage disequilibrium (LD). As a running example, we consider identifying allelic series for circulating lipid traits, drawing on data from the UK Biobank, Million Veterans Program, and Trans-Omics of Precision Medicine Program. Through extensive analyses of real and simulated data, we demonstrate that COAST-SS provides p-values effectively equivalent to those from the original COAST. Interestingly, we find that when LD is low, as is expected among rare variants, COAST-SS is robust to misspecification of the LD matrix, providing valid inference even when the LD matrix is set to the identity matrix. We explore several strategies for annotating the pathogenicity of variants supplied to COAST-SS, finding that they often yield similar power for detecting candidate allelic series. Lastly, we employ COAST-SS to screen for lipid trait allelic series in a meta-analyzed cohort of up to 840K subjects. COAST-SS has been incorporated into the publicly available `AllelicSeries R` package.

## Introduction

An allelic series is a collection of mutations in a gene or pathway that leads to a spectrum of possible phenotypes [1]. Allelic series that modulate gene activity or functionality provide the opportunity to characterize a gene’s dose-response behavior from the natural genetic variation in a population [2]. Such genes make appealing therapeutic targets because the impact of pharmacological modulation can be predicted from the impacts of observed genetic alterations [2]. The natural range of functionality observed in the population can also shed light on a candidate modulator’s therapeutic window and safety profile [3]. Moreover, with the rise of gene editing technologies, understanding allelic series could guide genetic interventions aimed at achieving beneficial phenotypic changes [4]. Among mutations that affect gene functionality, rare variants can have larger effect sizes than common variants, as purifying selection prevents large-effect deleterious variants from becoming common [5]. Thus, rare variants are particularly informative for characterizing allelic series.

The coding-variant allelic series test (COAST) is a rare-variant association test tailored for the identification of genes harboring allelic series [6]. COAST operates on three classes of coding variants, as annotated by the Ensembl Variant Effect Predictor (VEP) [7]: benign missense variants (BMVs), deleterious missense variants (DMVs), and protein truncating variants (PTVs). By means of an adjustable sequence of weights, which specifies the pattern of effect size sought, COAST targets genes where the impact on the phenotype increases monotonically with mutational severity. COAST was the first rare-variant association test specifically designed not simply to determine whether a gene is associated with a phenotype, but whether that association constitutes a dose-response relationship. However, the original implementation of COAST requires individual-level data [6], which is restrictive insofar as access to individual-level data is often limited by privacy concerns or logistical hurdles. In contrast, single-variant summary statistics from genome-wide association studies (GWAS) are often publicly available, even to relatively low minor allele frequencies. To facilitate their application, many methods originally developed in the context of individual-level data have been adapted to accept GWAS summary statistics as input, including heritability estimation, fine-mapping, Mendelian Randomization, and polygenic risk prediction [8, 9]. Doing so not only democratizes research, expanding the group of researchers who can analyze and derive insights from a data set, but also generally accelerates computation, benefiting even those who have access to the original data [10].

Here we extend COAST to enable estimation and inference starting from commonly available GWAS summary statistics. Specifically, COAST-SS performs allelic series testing starting from per-variant effect sizes and standard errors, along with estimates of the minor allele frequency (MAF) and local linkage disequilibrium (LD). We first validate that COAST-SS, working from summary statistics, can recover essentially the same p-value as the original COAST, working from individual-level data. Our example phenotypes throughout are a collection of 4 commonly-studied lipid traits: total cholesterol, high-density lipoprotein (HDL), low-density lipoprotein (LDL), and triglycerides. We proceed to evaluate the operating characteristics of COAST-SS, including type I error and power, as applied to both real and simulated data. Of the inputs required by COAST-SS, the rare variant LD matrix is the least likely to be available. We examine the performance of COAST-SS when the provided LD matrix is either approximate or absent altogether. We find that COAST-SS is robust to the LD matrix, providing valid inference even when supplied with the identity matrix. Compared with other rare-variant association tests, a unique aspect of COAST(-SS) is the classification of variants into 3 distinct categories (BMVs, DMVs, PTVs), across which the effect on the phenotype is expected to increase. We explore the impact on power of alternative schemes for weighting or partitioning variants into pathogenicity categories, using scores from AlphaMissense [11], ESM1b [12], MetaRNN [13], and PrimateAI [14]. Results are generally stable across the annotation and weighting schemes explored thus far. Lastly, we meta-analyze lipid trait summary statistics from the UKB and Million Veterans Program (MVP) to achieve a sample size of up to 840K for allelic series discovery. Associations from this analysis are validated with supporting evidence from multiple sources, including Genebass [15], the NHGRI-EBI GWAS Catalog [16], and the NHLBI Trans-Omics for Precision Medicine (TOPMed) Program [17].

## Material and Methods

### Recap of COAST

Let *Y* denote a quantitative phenotype (e.g. circulating lipid trait), and *X* a set of covariates (e.g. age, sex, ancestry), including an intercept. Let *G* = (*G*_1_, …, *G*_*J*_), with *G*_*j*_ ∈ {0, 1, 2}, denote the genotype at the *J* rare coding variants within a given gene. We restrict to variants belonging to one of the following annotation categories: benign missense variants (BMVs), deleterious missense variants (DMVs), and protein truncating variants (PTVs). For the *j*th variant, let *A*_*j*_ = 1 denote a BMV, *A*_*j*_ = 2 a DMV, and *A*_*j*_ = 3 a PTV. Denote by *N* = (*N*_1_, *N*_2_, *N*_3_) the total counts of BMVs, DMVs, and PTVs in the gene, respectively, and note that 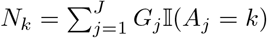 for *k* ∈ {1, 2, 3}. The **baseline allelic series model** is:

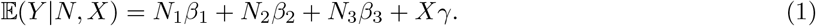

Given a set of **allelic series weights** *w* = (*w*_1_, *w*_2_, *w*_3_), the **allelic series sum model** is:

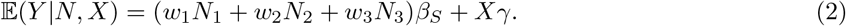

Note that the absolute scale of the allelic series weights (*w*_1_, *w*_2_, *w*_3_) is arbitrary, as scaling all the weights by a factor of *ω* ≠ 0 while scaling *β*_*S*_ by a factor of *ω*^*−*1^ would result in the same model. Without loss of generality, in this work, we scale the weights to lie in the [0, 1] interval, with *w*_3_ fixed at 1. Building on the sequence-kernel association test (SKAT) [18], the **allelic series SKAT model** is:

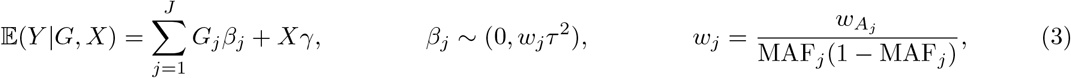

where 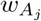 is the allelic series weight corresponding to the *j*th variant’s annotation, and MAF_*j*_ is the variant’s minor allele frequency. The overall **omnibus test statistic** is calculated by aggregating the p-values of the *M* = 3 component tests using the Cauchy combination method [19]:

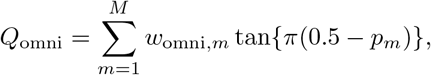

where the omnibus weights (*w*_omni,*m*_) are selected to give burden- and SKAT-type tests equal influence. The null hypothesis of the omnibus test is that *β*_1_ = *β*_2_ = *β*_3_ = 0 in (1), *β*_*S*_ = 0 in (2), and *τ* ^2^ = 0 in (3). Rejection of *H*_0_ by any of these tests encourages the omnibus test to reject. The final COAST p-value is calculated via:

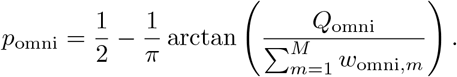

### Discrepancies from the original COAST

The original COAST [6] included an **allelic series max model**:

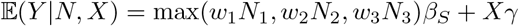

intended to capture cases where, for example, the presence of a loss-of-function PTV renders the presence of additional less-damaging variants moot. The allelic series max model is omitted from COAST-SS as it cannot be calculated from standard summary statistics. In addition, the original COAST performed burden testing with both count (additive) and indicator (dominant) genotype encodings. This distinction is dropped in COAST-SS as users of summary statistics typically do not have control over the genotype encoding, and the additive model is used most frequently in practice.

### COAST from Summary Statistics

#### Baseline model from summary statistics

For exposition, the models in the previous section were subject-level. Hereafter, we adopt matrix notation, with subjects as rows. Suppose we have *n* independent subjects. The baseline allelic series model is expressible as:

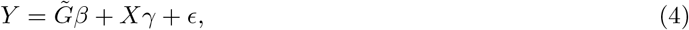

where 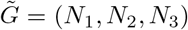 is an *n ×* 3 matrix, *β* = (*β*_1_, *β*_2_, *β*_3_) is a 3 *×* 1 coefficient vector, and *ϵ ~ N* (0, *σ*^2^*I*) is an *n ×* 1 vector of residuals. Note that the normality assumption on the residuals can be relaxed to the requirements that 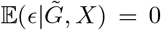 and 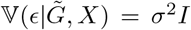 Define the *n × n* orthogonal projection matrix: *Q*_*X*_ = *I − X*(*X*^*′*^*X*)^*−*1^*X*^*′*^. The 3 *×* 1 score equation for *β* is:

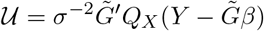

Under the *H*_0_ : *β* = 0, the score reduces to 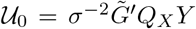, whose 3 *×* 3 variance-covariance matrix is 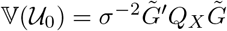. The *H*_0_ : *β* = 0 could be evaluated via:

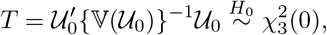

however this test requires access to the individual level data. We next develop an approximate test that can be evaluated using only summary statistics.

Suppose we have access to summary statistics from the single variant association models:

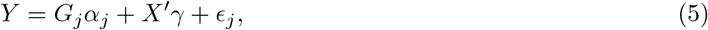

specifically 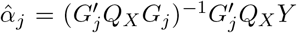 and 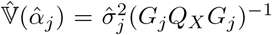. Note that 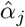 and 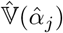 are scalars. Since under 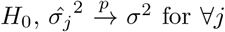, the sampling variance estimate 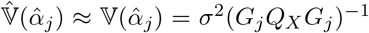. Let 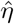 denote the vector of per-variant score statistics:

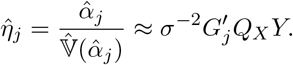

Define the *J ×* 3 variant category indicator matrix *D* such that:

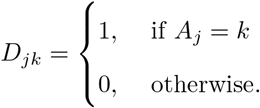

Then, 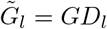 where *D*_*l*_ is the *l*-th column of *D*, and thus 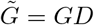. That is:

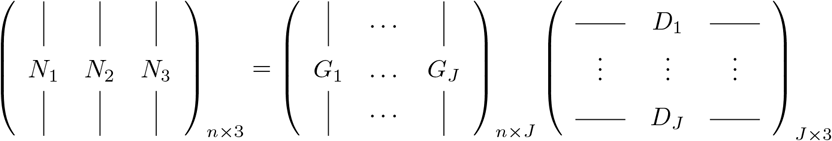

Now, we can express the score statistic above and its variance by:

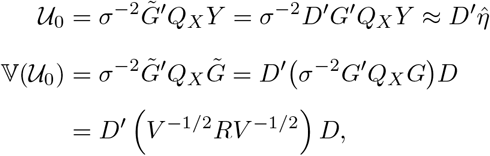

where the 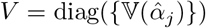 is the *J × J* matrix of per-variant sampling variances, which can be approximated by 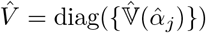, and the *j, k*-th element of the *J × J* correlation matrix *R* is:

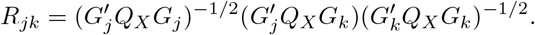

If we assume that the covariates are at most mildly correlated with the genotypes (which is reasonable for rare variants in an ancestrally homogeneous population) and are included in the model mainly for improving precision, *R*_*jk*_ can be approximated by the LD between the variants *j* and *k*. Hence, we can use the estimated LD matrix 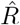 to replace *R* in the above equations to arrive at 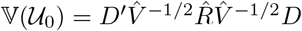. Overall, the test statistic is:

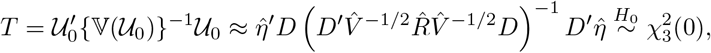

which can be calculated entirely from summary statistics.

#### Sum model from summary statistics

The allelic series sum model is a constrained case of the baseline model where *β*_*l*_ = *w*_*l*_*β*_*S*_, *w*_*l*_ is the allelic series weight for variant category *l*, and *β*_*S*_ is a common scalar. We can algebraically express this model as:

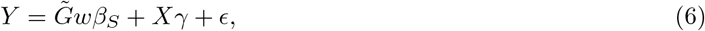

where *w* is the 3 *×* 1 allelic series vector (*w*_1_, *w*_2_, *w*_3_) and *β*_*S*_ is a scalar. Hence, the score statistic and its variance (now scalars) are given by:

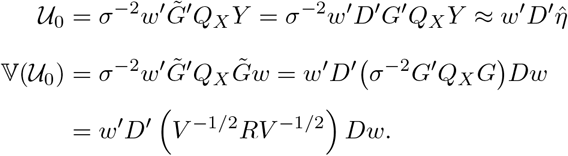

Therefore, the test statistic for the allelic series sum model is:

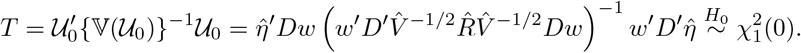

#### SKAT model from summary statistics

Recall that, under *H*_0_ : *β* = 0, the score statistic from the joint model (4) is 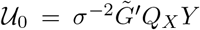. The score statistic for the SKAT model in (3) takes a similar form, *𝒱* = *σ*^*−*2^*G*^*′*^*Q*_*X*_ *Y*, except *G* = (*G*_1_, …, *G*_*J*_) is now the individual-variant genotype matrix. The SKAT statistic [18] is a variance component test of *H*_0_ : *τ* ^2^ = 0 in (3) based on the quadratic form *T* = *𝒱*^*′*^*K𝒱*, where *K* is a symmetric, positive definite kernel matrix. The kernel matrix *K* is calculated as a known function of the MAFs of the individual variants. The distribution of *T* is a mixture of 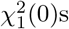s, whose mixing proportions *λ*_*j*_ for *j* ∈ {1, …, *J*} are the eigenvalues of 𝕍^1*/*2^(*𝒱*)*K*𝕍^1*/*2^(*𝒱*). From the variant-level summary statistics 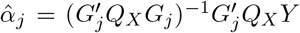 and 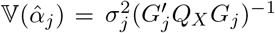 we calculate the entries of the score vector by 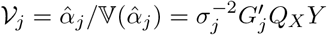. The covariance of the per-variant scores is 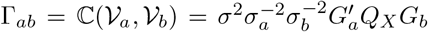. Writing 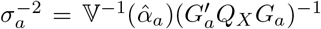 and 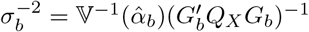, the covariance becomes:

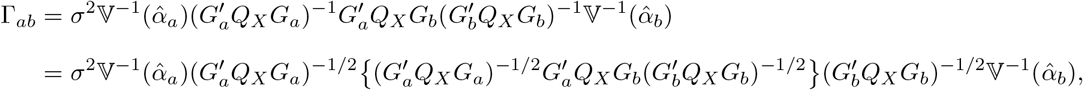

which we approximate by 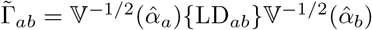. Having calculated *𝒱* and estimated its covariance matrix Γ, we calculate the Cholesky decomposition of Γ = *LL*^*′*^, then take the spectral decomposition of *L*^*′*^*KL* to obtain the set of eigenvalues *λ*_*j*_ *> ϵ* = 10^*−*8^. The *H*_0_ : *τ* ^2^ = 0 is evaluated by comparing the test statistic *T* = *𝒱*^*′*^*K𝒱* to the reference distribution 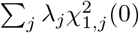. The final p-value is calculated via Davies’ algorithm, as implemented by the `CompQuadForm` package (v1.4.3) [20] in `R`.

### Simulation Studies

#### Type I error

Type I error studies were performed using real genotypes and phenotypes from the same cohort of 150K unrelated subjects examined in the original COAST publication [6]. The phenotype was permuted LDL. Genotype preparation is described in detail below. In total, more than 500 genome-wide rare-variant association studies (RVAS) were performed, each including around 17K genes, resulting in more than 10M realizations of the COAST-SS p-value under the null.

#### Power

In order to have fine control over the genetic architecture, power studies were based on simulated data, using the framework described previously [6]. Each simulated gene contained 10^2^ variants. The expected frequencies of BMVs, DMVs, and PTVs were 50%, 40%, and 10%, mirroring their empirical frequencies in the human genome. A random subset of variants, between 25% and 100%, was selected as causal. The effect sizes of non-causal variants were set to zero. Burden phenotypes were simulated from model (1) with various settings for (*β*_1_, *β*_2_, *β*_3_). For SKAT phenotypes, per-variant effect sizes were generated as: 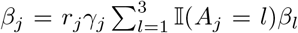, where *r*_*j*_ ∈ {*−*1, 1} is a random sign, *γ*_*j*_ *~* Γ(1, 1) is a positive random scalar with 𝔼(*γ*_*j*_) = 1 and 𝕍(*γ*_*j*_) = 1, and the term 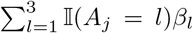 incorporates an additional fixed weight depending on the variant’s annotation category.

### Preparation of UK Biobank Data

#### Whole exome sequencing data

Variants in the full UKB WES variant callset (*N* = 469K, [21]) were removed if they were located *>* 100 bp outside the exome capture regions, within the ENCODE blacklist [22], or within low-complexity regions [23]. We further filtered to retain only autosomal, biallelic variants meeting the following criteria: allele balance fractions of ≤ 0.1 (homozygous) or ≥ 0.2 (heterozygous), call rate ≥ 0.95, mean depth (DP) 12 ≤ DP *<* 200, mean genotype quality GQ ≥ 20, Hardy-Weinberg equilibrium *p >* 1*×*10^*−*10^, and inbreeding coefficient *F > −*0.3. Variants were annotated with VEP (v109.3) [7] using the following transcript consequences (for any transcript of a gene): protein-truncating variants (PTV): `splice acceptor variant`, `splice donor variant`, `stop gained`, or `frameshift variant`; missense: `missense variant`, `inframe deletion`, `inframe insertion`, `stop lost`, `start lost`, or `protein altering variant`. Missense variants were annotated to be damaging (DMV) if predicted to be `probably damaging` or `possibly damaging` by PolyPhen [24] and `deleterious` or `deleterious low confidence` by SIFT [25]. Missense variants were annotated to be benign (BMV) if predicted to be `benign` by PolyPhen and `tolerated` or `tolerated low confidence` by SIFT [26, 27]. PTVs were further filtered to remove low confidence loss-of-function by LOFTEE [28]. The most severe consequence per variant-gene pair was retained.

#### Quality control

Sample-level quality control (QC) was performed using 88,813 high-confidence variants passing the above filtering criteria, which were LD-pruned to *r*^2^ *<* 0.05. To mitigate confounding due to population structure, samples were first filtered to individuals without sex chromosome aneuploidy, within *±*7 standard deviations of the mean of the first six genotype principal components [29], self-reported ethnic background of `White British`, `Irish`, or `White`, had call rate ≥ 0.95, mean DP ≥ 19, and mean GQ ≥ 47.5. We regressed out the top 20 genetic principal components from the following sample QC metrics and removed samples that were *>* 6 median absolute deviations from the median for any of the following metrics: number of SNPs, number of heterozygous variants, number of homozygous alternate variants, number of insertions, number of deletions, number of transitions, number of transversions, the ratio between the number of transitions and transversions, the ratio between the number of heterozygous and homozygous alternate variants, and the ratio between the number of insertions and deletions [28]. The sample size after filtering was 390,865. All data preprocessing was performed in `Hail v0.2.116` [30].

#### UKB cohorts

For analyses where comparison with the original COAST was desired, we used the same subset of 145,735 subjects as in [6]. We refer to this subset as the 150K cohort. Type I error simulations were also performed within the 150K cohort because the operating characteristics of statistical methods typically improve with increasing sample size, making the evaluating within the smaller cohort more challenging. The remaining UKB-based analyses were performed within the superset of 390,865 subjects, referred to as the 390K cohort.

#### Association analysis

Processed WES data were restricted to BMVs, DMVs, and PTVs with a sample MAF ≤ 1% unless otherwise stated. Following [19, 31], ultra-rare variants, those with a sample MAC ≤ 10, were collapsed into a single aggregated variant separately for each gene *×* variant category. Genes were required to have at least 3 distinct rare variants (including the aggregated variants) for inclusion in the genome-wide screen. Lipid traits were standardized via the rank-based inverse normal transformation [32]. Per-variant effect sizes and standard errors were calculated among 390,865 individuals adjusting for a sparse genetic relatedness matrix in `Saige` (`v1.3.0`) [33]. MAFs and per-gene LD matrices were calculated in `numpy` (`v1.23.5`). COAST-SS was performed using the `AllelicSeries R` package (`v1.1.2`) with full in-sample LD matrices.

### Meta-analysis

Lipid trait summary statistics for 449,042 subjects of European ancestry from the Million Veterans Program (MVP) [34] were downloaded from the database of Genotypes and Phenotypes (dbGaP), accession number phs001672.v11.p1, and filtered to those with a MAF ≤ 0.1%. A stricter MAF threshold was applied here as LD information was unavailable for MVP. The MVP summary statistics were combined with those from the UKB 390K cohort via inverse-variance-weighted (i.e. fixed-effects) meta-analysis. The meta-analysis was performed via the `R meta` package (`v8.0`) [35]. Variants present in only one cohort were retained, with the effect sizes and standard errors unchanged. COAST-SS was run borrowing approximate LD information from the UKB where available.

### Overlap Analysis

For each lipid trait, common-variant associations were downloaded from the NHGRI-EBI GWAS Portal [16] on 2024-10-21 and filtered to those with *p <* 5 *×* 10^*−*8^. Rare-variant associations from both putative loss of function (pLoF) variants and missense variants, including low-confidence pLoF variants and in-frame insertions or deletions, were obtained from the Genebass Portal [15]. Associations were filtered to those Bonferroni significant based on the number of genes with non-missing p-values, and the union of significant associations from the pLoF and missense analyses was taken.

#### TOPMed analysis

As an additional validation in a multi-ancestry setting, we applied COAST-SS to lipid trait summary statistics from Freeze 8 of the NHLBI TOPMed Program [17], obtained through dbGaP. This cohort is composed of 66,329 participants from diverse ancestry groups, including 44.48% European, 25.60% Black, 21.02% Hispanic, 7.11% Asian, and 1.78% Samoan. Summary statistics were generated from a pooled multi-ancestry mixed-model analysis, with adjustment for ancestry-related covariates, including genetic PCs and study subgroups [17]. Summary statistics were only available for variants with a sample MAF *>* 0.1%, and subsequent allelic series analysis was restricted to variants with a MAF ≤ 1%. Although imperfect given the multi-ancestry composition of TOPMed, COAST-SS was run borrowing LD information from the UKB where available. Significant gene-trait associations from allelic series analysis of the meta-analyzed UKB+MVP cohort were queried for replication in TOPMed at Bonferroni and nominal significance.

## Results

### COAST-SS runs COAST starting from summary statistics

As detailed in the Methods, COAST-SS performs a rare coding-variant allelic series test starting from summary statistics. The inputs to COAST-SS are the per-variant effect sizes (i.e. 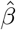s) and standard errors (SEs), together with per-variant MAFs and the local variant-by-variant LD matrix. We first validated that COAST-SS can recover essentially the same p-value as the original COAST when starting from summary statistics. The probability-probability (PP) plots in **Figure 1** compare the p-values from COAST-SS with those from COAST for 4 lipid traits (cholesterol, HDL, LDL, triglycerides) using genotypes from 17K genes and phenotypes from 150K subjects in the UKB. Exact agreement is not expected because COAST includes component tests that could not be computed from standard summary statistics. Nonetheless, the *R*^2^ is at least 0.999 in all cases, and the estimated slopes are within 2% of the identity. Although its close agreement with the original COAST suggests that COAST-SS likewise provides a valid test of association, we confirmed this by performing extensive type I error studies. Specifically, RVAS with real genotypes were performed on random permutations of LDL cholesterol, generating 10M *p*-values under the null. The quantile-quantile plots in **Figure S1** demonstrate that p-values of COAST-SS, as well as its 3 component tests (the allelic series baseline, sum, and SKAT models), are uniformly distributed in the absence of association. **Table 1** provides estimates of the type I error at *α* = 2.5 *×* 10^*−*6^, the Bonferroni threshold when adjusting for 20K genes, as well as the expected *χ*^2^ statistic and genome-inflation factor *λ*_*p*_ at the center (*p* = 0.5) and in the lower tail (*p* = 0.001) of the p-value distribution. In all cases, the type I error is at or below the nominal threshold, while the expected *χ*^2^ statistics and genomic-inflation factors are near the expected value of 1. COAST-SS exhibits some deflation in the center of its p-value distribution, an expected consequence of employing the Cauchy combination [36], however this is immaterial insofar as only p-values in the lower tail are of interest. Overall, we find the COAST-SS provides a valid test of association.

**Table 1:** COAST-SS controls type I error under the null. Results are based on 10M p-values generated using real data on 150K subjects from the UKB. The phenotype was LDL, which was permuted to break any association with genotype. Type I error (×10^6^) is reported at *α* = 2.5 ×10^*−*6^, which is the Bonferroni threshold when adjusting for 20K genes; the expected value is 2.5. Also reported are the expected *χ*^2^, the median genomic inflation factor *λ*_0.5_, and the tail genomic inflation factor *λ*_0.001_. LD method refers to how the LD matrix was estimated; original means the test was performed with the original COAST (i.e. using individual-level data).

| Test | LD Method | T1E ($\times 10^6$) | $\mathbb{E}(\chi^2)$ | $\lambda_{0.5}$ | $\lambda_{0.001}$ |
| --- | --- | --- | --- | --- | --- |
| Baseline | Original | 2.10 | 1.00 | 1.00 | 1.00 |
| Baseline | Full LD | 2.10 | 1.00 | 1.00 | 1.00 |
| Baseline | Sub-sample LD | 2.40 | 1.00 | 1.00 | 1.00 |
| Baseline | Identity | 2.30 | 1.00 | 1.00 | 1.00 |
| Allelic Sum | Original | 1.80 | 1.00 | 1.00 | 1.00 |
| Allelic Sum | Full LD | 1.80 | 1.00 | 1.00 | 1.00 |
| Allelic Sum | Sub-sample LD | 2.00 | 1.00 | 1.00 | 1.00 |
| Allelic Sum | Identity | 2.40 | 1.01 | 1.01 | 1.01 |
| Allelic SKAT | Original | 1.50 | 1.00 | 0.98 | 1.00 |
| Allelic SKAT | Full LD | 2.00 | 1.00 | 0.99 | 1.00 |
| Allelic SKAT | Sub-sample LD | 2.20 | 1.00 | 0.99 | 1.00 |
| Allelic SKAT | Identity | 2.40 | 1.01 | 0.99 | 1.01 |
| COAST-SS | Original | 2.40 | 1.00 | 0.89 | 1.00 |
| COAST-SS | Full LD | 2.00 | 1.00 | 0.90 | 1.00 |
| COAST-SS | Sub-sample LD | 2.20 | 1.00 | 0.90 | 1.01 |
| COAST-SS | Identity | 2.50 | 1.01 | 0.90 | 1.01 |

**Figure 1:**
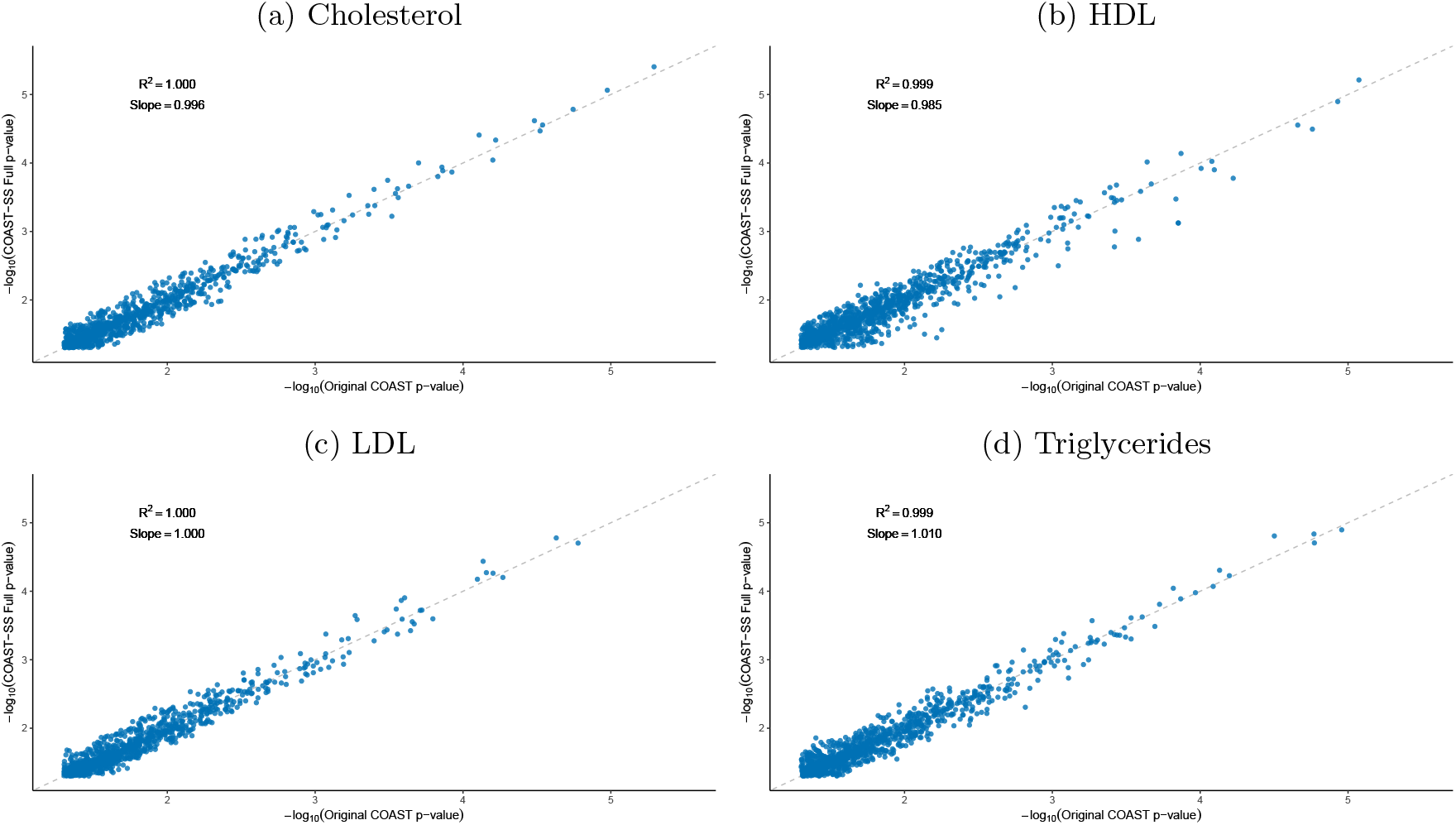
COAST from summary statistics (COAST-SS) recovers essentially the same p-value as the original COAST. Note that perfect agreement is not expected because COAST-SS includes a different set of component tests than COAST. Shown are genes with a *P* ≤ 0.05 for 4 lipid traits from the UKB. The sample size was *N* = 150K. The dotted gray line is the identity. The ideal value for both the *R*^2^ and the estimated slope is 1.

### COAST-SS remains valid in the absence of perfect LD information

Among the summary statistics required by COAST-SS, the rare-variant LD matrix is the one least readily available. We evaluated several methods of calculating or approximating the LD matrix supplied to COAST-SS. The gold-standard approach utilizes the full sample in which the association statistics were generated to calculate the LD. The sub-sample approach utilizes a random sample of 50K unrelated subjects drawn with replacement, emulating a case where the LD matrix is calculated via a different sample from the same population. As rare variants are, by definition, present in only a small proportion of subjects, some variants present in the full sample will not appear in any given sub-sample. In these cases, the corresponding row and column of the sub-sample LD matrix were taken from the identity matrix. The final approximation, emulating a case where LD information is unavailable, simply supplies an identity matrix to COAST-SS. Although ignoring correlations between variants would likely compromise a test’s operating characteristics in the common variant setting, in the rare variant setting the LD between variants is often negligible [37]. Indeed, across all 17K genes in our RVAS studies, the mean between-variant *R*^2^ in the full sample LD matrices was 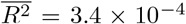, and for half of all genes the mean 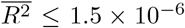. As a comparator for COAST-SS performed with the full-sample, sub-sample, and identity LD matrices, we present the calculation performed with the original COAST, using individual-level data. Interestingly, the type I error studies in **Table 1** and **Figure S1** indicate that all LD methods, including the identity matrix, maintain control of the type I error. Note that these experiments utilized real genotypes, with their inherent LD structure. Moreover, the PP-plots in **Figure S2** exhibit tight concordance between applying COAST-SS with full LD versus the identity matrix to real-data analyses of 4 lipid traits in an expanded cohort of 390K subjects from the UKB. Across phenotypes simulated from a variety of genetic architectures, all methods supplied nearly identical power (**Figures 2**, **S3**). These results suggest that COAST-SS is robust to the LD matrix, and may even be run with an identity matrix, provided it is reasonable to assume the LD among the supplied variants is small. Out of an abundance of caution, subsequent sections continue to run COAST-SS with the full ins-sample LD matrix unless otherwise stated.

**Figure 2:**
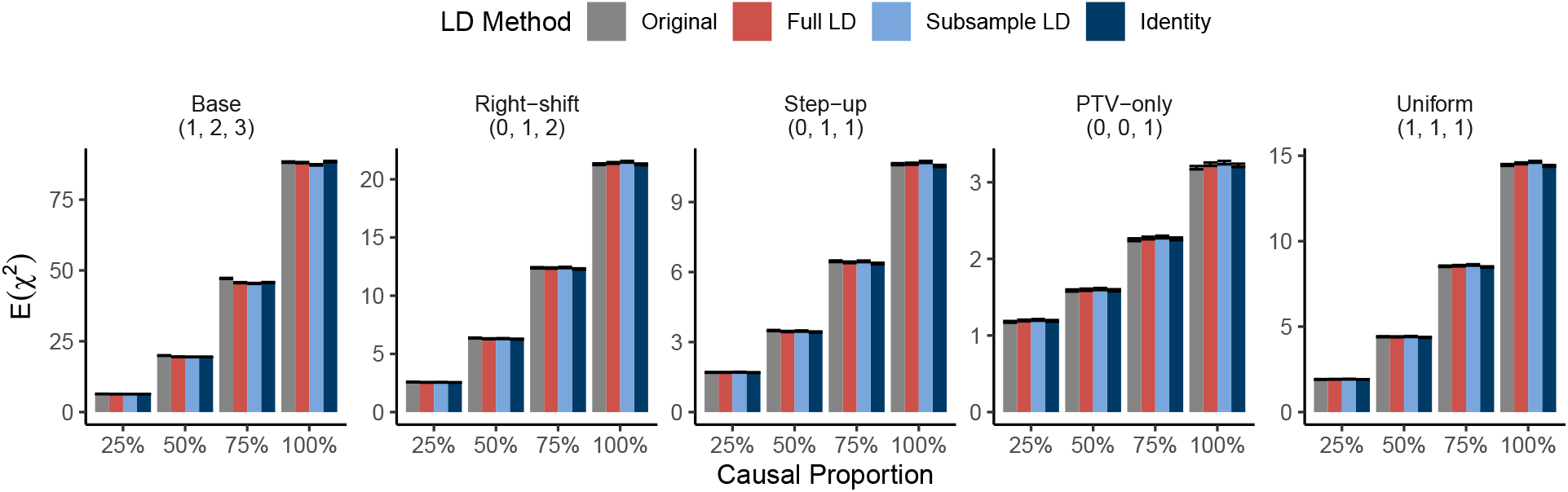
COAST-SS has equivalent power to COAST, and is robust to the provided LD matrix. Original refers to COAST run with individual-level data; full LD was calculated using all 150K subjects; sub-sampled LD was calculated using a random sample of 50K subjects, drawn with replacement; identity ignores LD. Results are presented under 5 different generative models. The tuple above each set of results indicates the true effect sizes of BMVs, DMVs, and PTVs in the generative model. Power is quantified by the expected *χ*^2^ statistic. The simulation sample size was *N* = 10^4^. Error bars are 95% confidence intervals for the mean across *R* = 10^5^ simulation replicates.

### COAST-SS substantially accelerates genome-wide analyses

A central motivation for the development of COAST-SS was to enable allelic series testing when individual-level data are unavailable. As shown in **Table 2**, a secondary advantage is that COAST-SS dramatically reduces the time required to perform a genome-wide analysis. Compared with the original COAST, COAST-SS reduces the runtime by approximately 100*×*. With COAST-SS, the expected time to complete a 20K genes scan is 113 ms *×* 20000 = 38 mins, which we further reduced by running batches of genes in parallel. There was not an appreciable difference in runtime between COAST-SS with a dense LD matrix versus the identity matrix. Of course, the reason COAST-SS is substantially faster is that much of the computation has been performed prior to allelic series testing. The computation of per-gene LD matrices, per-variant MAFs, and per-variant annotations is a one-time cost, and the resulting variant-level information can be reused for analyzing any subsequent phenotype. We performed the LD and MAF calculations with `Hail` [30] and `Python`, and annotation using Ensembl’s `VEP` [7]. The other prerequisite to running COAST-SS is calculating the per-variant effect sizes and SEs. Many highly optimized tools are available for this task, including `PLINK` [38], `GCTA` [39], `BOLT-LMM` [40], `Saige` [33], `Regenie` [41], and `STAARpipeline` [42].

**Table 2:** COAST-SS substantially accelerates genome-wide analyses. 5 genome-wide RVAS were conducted using real data on 150K subjects from the UKB. Results refer to per-mean runtimes and are measured in milisections (ms). Min refers to the minimum, *p*_25_ the 25th percentile, *p*_50_ the median, *µ* the mean, *p*_75_ the 75th percentile, Max the maximum.

| Test | LD Method | Min | $p_{25}$ | $p_{50}$ | $\mu$ | $p_{75}$ | Max |
| --- | --- | --- | --- | --- | --- | --- | --- |
| COAST | Original | 7007.00 | 9822.00 | 10309.00 | 10653.14 | 11062.00 | 62635.00 |
| COAST-SS | Full LD | 3.00 | 68.00 | 79.00 | 113.21 | 99.00 | 2319.00 |
| COAST-SS | Identity | 2.00 | 69.00 | 80.00 | 113.88 | 102.00 | 1834.00 |

### Alternate annotation and weighting schemes provide consistent results

As described in the whole exome sequencing methods, a baseline annotation of BMV, DMV, or PTV was assigned to each variant on the basis of transcript consequence information from VEP [7] and qualitative missense pathogenicity classifications from SIFT [25] and PolyPhen-2 [24]. To explore whether incorporating information from recent variant effect prediction models could improve power for allelic series detection, quantitative pathogenicity scores from several models, including AlphaMissense [11], ESM1b [12], MetaRNN [13], and PrimateAI [14], were queried from dbNSFP `v4` [43]. **Figure S4(a)** depicts the correlation among the quantitative pathogenicity scores, which was 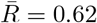 on average, and **Figure S4(b)** depicts the mean pathogenicity score within each of the baseline annotation categories. Two strategies for integrating the pathogenicity scores were explored. In the Annotation Experiment, all missense variants were reassigned to 1 of 3 categories based on tertiles of the pathogenicity score calculated genome-wide. PTVs, which are not scored by most missense pathogenicity models, were always assigned to category 3. Although categories defined in this way do not correspond precisely to “benign”, “damaging”, and “protein-truncating”, they nonetheless form a discrete, monotonic scale, across which the phenotypic impact of mutations is expected to increase. In the Weighting Experiment, each variant retained its baseline annotation category, but the allelic series weights were calculated, adaptively for each gene, as the mean value of the pathogenicity score for all variants within each category. When calculating these means, PTVs were assigned the maximum score of 1.

We applied COAST-SS to perform the Annotation and Weighting Experiments with summary statistics from 4 lipid traits (cholesterol, HDL, LDL, triglycerides) in the UKB 390K cohort. **Figure 3(a)** depicts the number of Bonferroni significant associations for each trait, and **Figure 3(b)** reports the mean *χ*^2^ statistic across Bonferroni significant genes. In general, the baseline annotation and weighting schemes were competitive with those informed by newer pathogenicity scores. Across annotation and weighting schemes, the p-values assigned to significant gene-trait pairs were highly concordant, with Spearman *R*^2^s exceeding 0.78 in all cases for the annotation experiment, and exceeding 0.98 in all cases for the weighting experiment (**Figure S5**). Most gene-trait associations detected under any annotation or weighting scheme were detected by all, and in cases where a gene-trait pair was detected by only some, the p-values tended to be suggestive where not significant. Comparing the annotation and weighting strategies, the latter approach, of calibrating the allelic series weights *w* = (*w*_1_, *w*_2_, *w*_3_) based on the mean pathogenicity scores within the baseline annotation categories, led to more Bonferroni significant associations on average (20.2 vs. 18.6). Given the consistency of findings across annotation and weighting schemes in real data, we also verified through simulations that informative annotations do in fact improve power. **Figure S6** confirms that, in cases where the expected phenotypic impact differs across annotation categories, providing the correct annotations increases power as compared with annotations that were either permuted or sampled uniformly at random.

**Figure 3:**
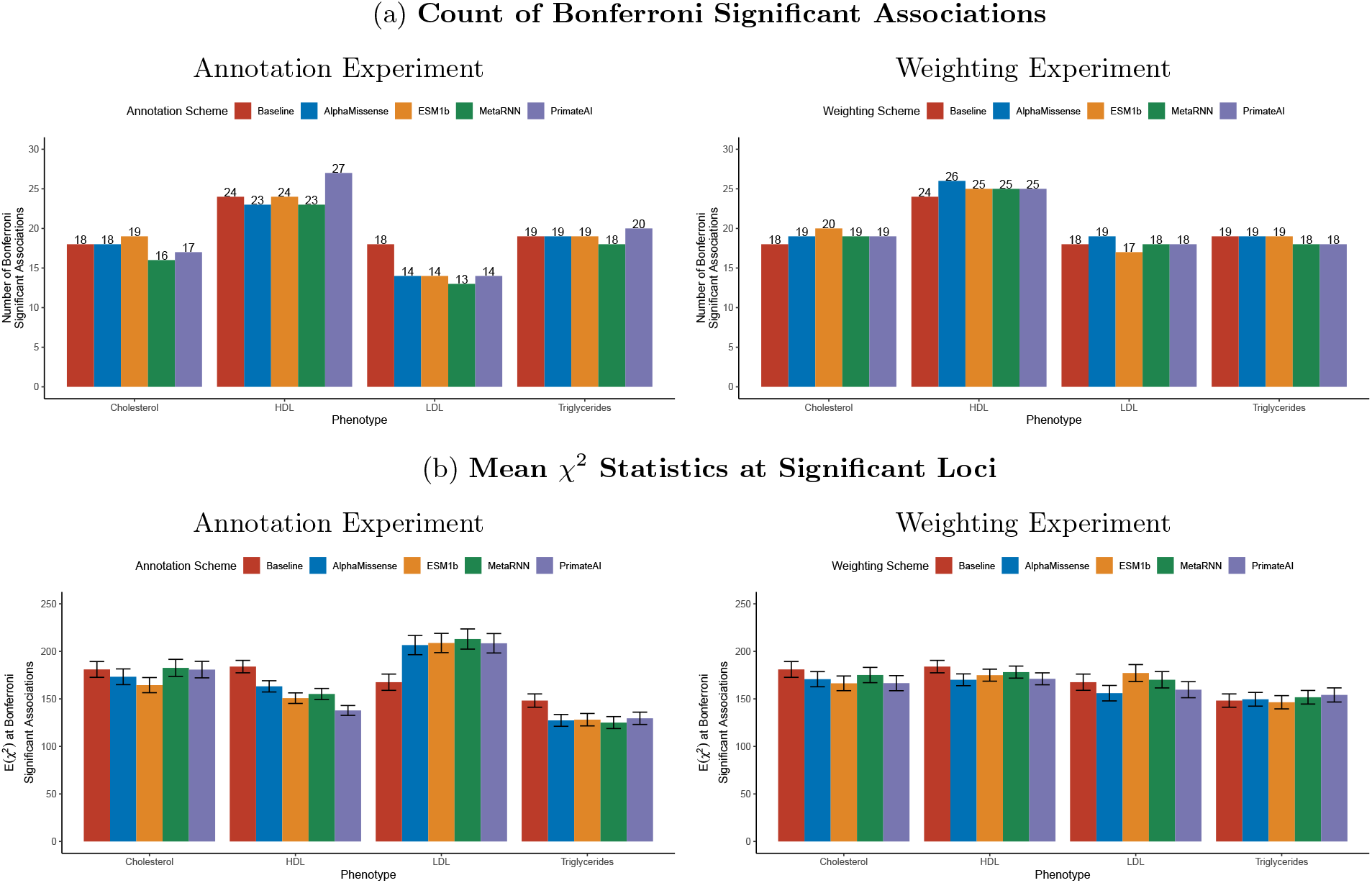
Different missense variant annotation schemes provided similar power for detecting associations with lipid traits. COAST-SS with full LD was run on summary statistics for lipid traits from 390K subjects in the UKB. Baseline is the original annotation scheme, which classifies variants into BMVs, DMVs, and PTVs. In the Annotation Experiment, missense variants were reassigned to 3 pathogenicity categories based on quantiles of a given score, while the weights were fixed at their default values, an evenly spaced increasing sequencing. PTVs were always assigned to the highest category. In the Weighting Experiment, the annotation scheme was fixed at baseline, but the allelic series weights were set, adaptively for each gene, to the mean value of the pathogenicity score for variants in that category. When taking the mean, PTVs were assigned the maximum weight of 1. (a) depicts the number of Bonferroni significant associations, adjusting for 17,295 tests of association, and (b) presents the mean *χ*^2^ statistics across the Bonferroni significant associations, a proxy for power. Error bars are 95% confidence intervals.

### COAST-SS identifies new candidate allelic series for lipid traits

Towards identifying new candidate allelic series for lipid traits, we meta-analyzed summary statistics from 450K subjects in the MVP [34] with those from the UKB 390K cohort, for a combined sample size of up to 840K. LD matrices were constructed by borrowing in-sample LD information from the UKB where possible, and assuming linkage equilibrium otherwise. The resulting Manhattan plots are presented in **Figure 4**. Two well-known lipid genes were identified as candidate allelic series for all 4 lipid traits: *APOB* and *ANGPTL3*. Apolipoprotein B (*APOB*) provides a scaffold for the transport and uptake of circulating lipids, while angiopoietin like 3 (*ANGPTL3*) regulates (inhibits) lipoprotein lipase activity [44]. In total, COAST-SS identified 108 Bonferroni significant associations: 21 for total cholesterol, 41 for HDL, 22 for LDL, and 24 for triglycerides. The Venn diagrams in **Figure S7** decompose the these counts according to whether the association was detected in the UKB 390K versus the combined UKB+MVP cohort. The majority of associations (77 of 79) detected in the UKB 390K cohort were also detected in the meta-analysis. Meta-analysis with MVP contributed an additional 31 associations (39.2% increase), of which 17 were with HDL. Upon meta-analysis, one gene (*UGT2B7*) dropped below the Bonferroni significance threshold, but remained suggestively significant, for both total cholesterol (UKB: *P* = 4.9 *×* 10^*−*7^, UKB+MVP: *P* = 5.4 *×* 10^*−*6^) and LDL (UKB: *P* = 7.9 *×* 10^*−*7^, UKB+MVP: *P* = 1.4 *×* 10^*−*5^).

**Figure 4:**
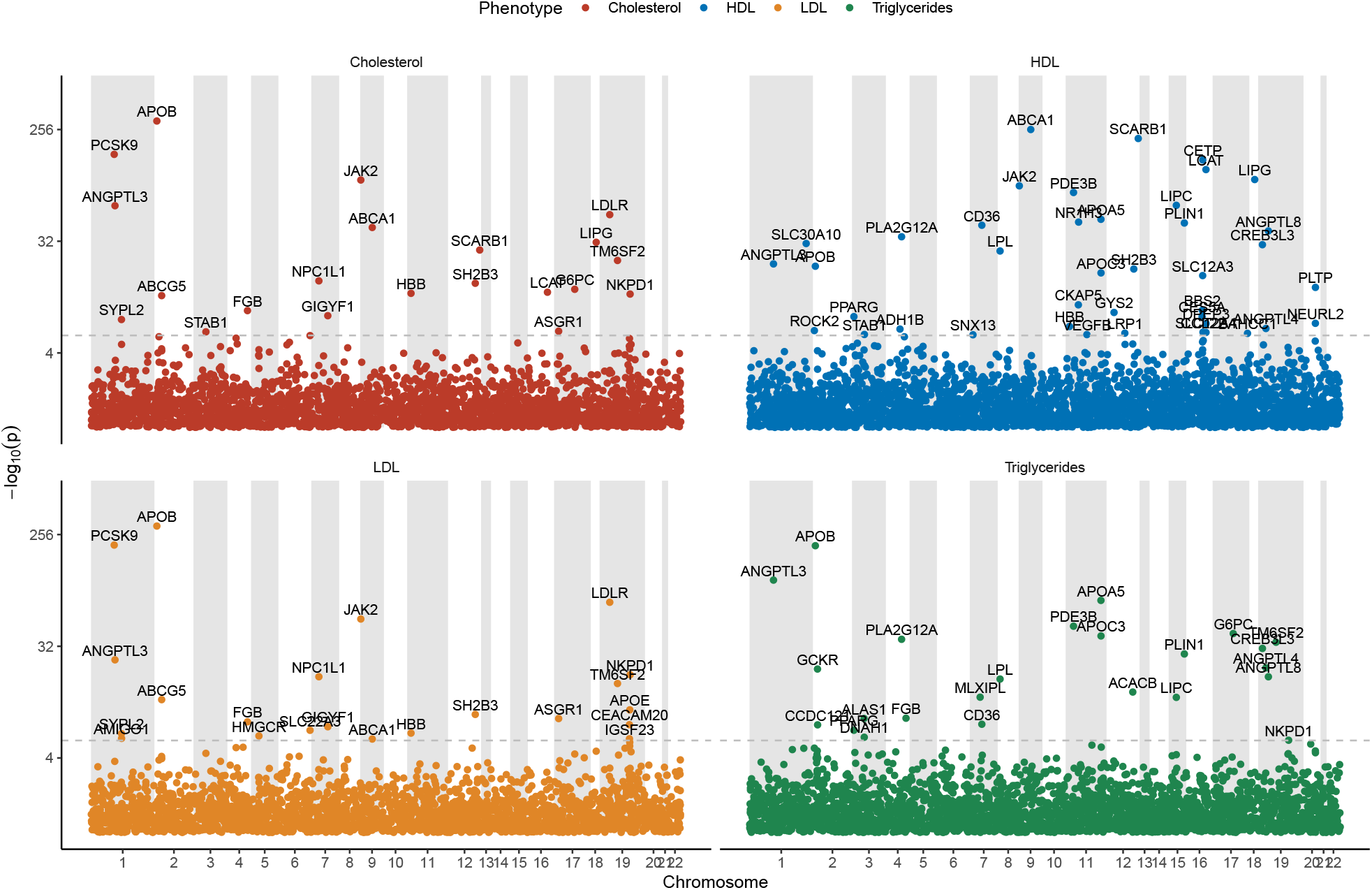
Manhattan plots for allelic series analysis of lipid trait summary statistics from the meta-analyzed UKB+MVP cohort. The analysis includes 17,503 genes. Gene symbols are shown for Bonferroni significant associations.

For each lipid trait, we overlapped the genes identified by COAST-SS in the meta-analyzed UKB+MVP cohort with previously reported associations from Genebass [15], to assess rare-variant support, and from the GWAS Catalog [16], to assess common-variant support. The results are presented in **Table 3(a)**. Among the 108 associations detected by COAST-SS, 80 had previous rare-variant support, while 28 did not. Of these, 23 had prior common-variant support, while 5 associations (all with HDL) are potentially novel. As seen in **Table 3(c)**, the significance of all 5 associations was driven by the introduction of data from the MVP. As an orthogonal validation, we applied COAST-SS to lipid trait summary statistics from 66K subjects in the multi-ancestry TOPMed cohort [17], and queried the findings from the UKB+MVP analysis for evidence of association in TOPMed. The results are shown in **Table 3(b)**. By comparison with Genebass or the GWAS Catalog, the replication rate in TOPMed was relatively modest (39.0% on average), which we attribute to the relatively small sample size size (*N* = 66K) coupled with the absence of summary statistics for variants with a MAF ≤ 0.1%, which selectively depletes larger effect size variants (e.g. PTVs and DMVs).

**Table 3:**
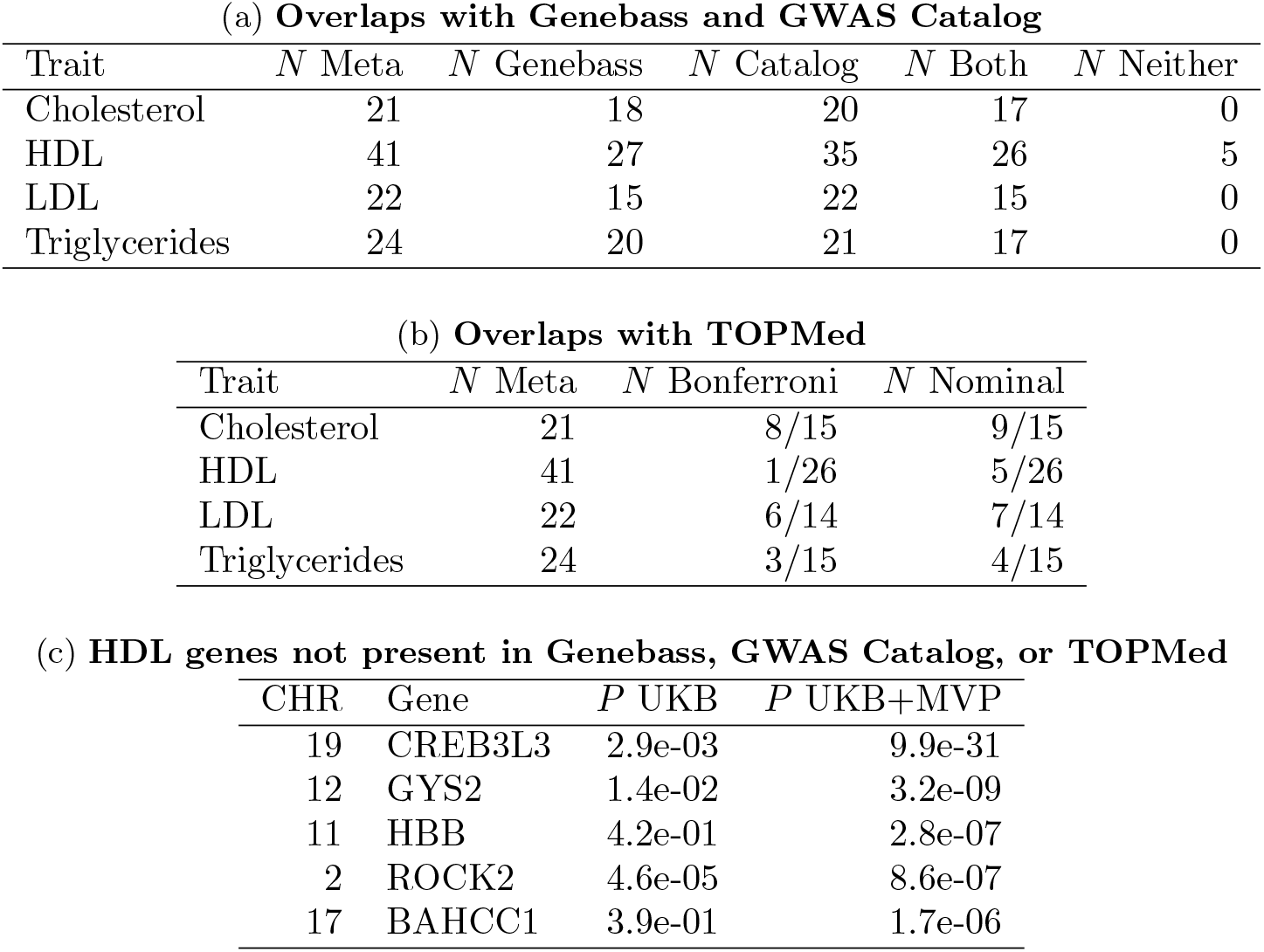
COAST-SS identifies potentially novel associations for HDL. In all cases, associations are Bonferroni significant unless otherwise noted. (a) Overlap of associations from COAST-SS applied to the UKB+MVP cohort with associations for the same traits from Genebass and the GWAS Catalog. *N* Meta is the total number of associations identified in the UKB+MVP analysis. *N* Catalog and *N* Genebass are the counts of associations also present in the GWAS Catalog and Genebass, respectively. *N* Both and *N* Neither are the counts of associations in both or neither the GWAS Catalog and Genebass. (b) Overlap of associations from the UKB+MVP analysis with associations from TOPMed at Bonferroni and nominal significance. The denominator shows the count of genes from the UKB+MVP that were present in the TOPMed analysis. (c) Genes associated with HDL in the UKB+MVP analysis not previously reported in Genebass or the GWAS Catalog, and not reaching significance in TOPMed.

| Trait | *N* Meta | *N* Genebass | *N* Catalog | *N* Both | *N* Neither |
| --- | --- | --- | --- | --- | --- |
| Cholesterol | 21 | 18 | 20 | 17 | 0 |
| HDL | 41 | 27 | 35 | 26 | 5 |
| LDL | 22 | 15 | 22 | 15 | 0 |
| Triglycerides | 24 | 20 | 21 | 17 | 0 |

(b) **Overlaps with TOPMed**
| Trait | *N* Meta | *N* Bonferroni | *N* Nominal |
| --- | --- | --- | --- |
| Cholesterol | 21 | 8/15 | 9/15 |
| HDL | 41 | 1/26 | 5/26 |
| LDL | 22 | 6/14 | 7/14 |
| Triglycerides | 24 | 3/15 | 4/15 |

Table 3: **COAST-SS identifies potentially novel associations for HDL.**
| CHR | Gene | *P* UKB | *P* UKB+MVP |
| --- | --- | --- | --- |
| 19 | CREB3L3 | 2.9e-03 | 9.9e-31 |
| 12 | GYS2 | 1.4e-02 | 3.2e-09 |
| 11 | HBB | 4.2e-01 | 2.8e-07 |
| 2 | ROCK2 | 4.6e-05 | 8.6e-07 |
| 17 | BAHCC1 | 3.9e-01 | 1.7e-06 |

**Figure 5** presents the mean absolute effect size by variant category for each of the 108 associations detected by COAST-SS. Among the 5 associations with HDL not previously reported by Genebass or the GWAS Catalog, 2 (*CREB3L3* and *HBB*) had a monotone increasing effect size pattern from BMVs to DMVs to PTVs. *CREB3L3* encodes a cyclic adenosine monophosphate (cAMP-)responsive transcription factor that regulates the expression of lipid metabolism genes [45]. Loss of function mutations in *CREB3L3* have been linked to hypertriglyceridemia [46, 47], likely explaining the association of this gene with triglycerides. In addition, *CREB3L3* inhibits *ANGPTL3* [48], which influences HDL levels through the regulation of lipoprotein lipase [49]. In Genebass, *ANGPTL3* is associated with all of HDL, LDL, triglycerides, and total cholesterol. *HBB* encodes the beta subunit of hemoglobin. Hemoglobin concentration has been associated with circulating lipoprotein size [50], and hemoglobin together with its scavenger haptoglobin have been localized to HDL particles in both murine models and coronary heart disease patients [51]. Common loss-of-function variants in *HBB* have previously been associated with both total cholesterol and LDL [52, 53].

**Figure 5:**
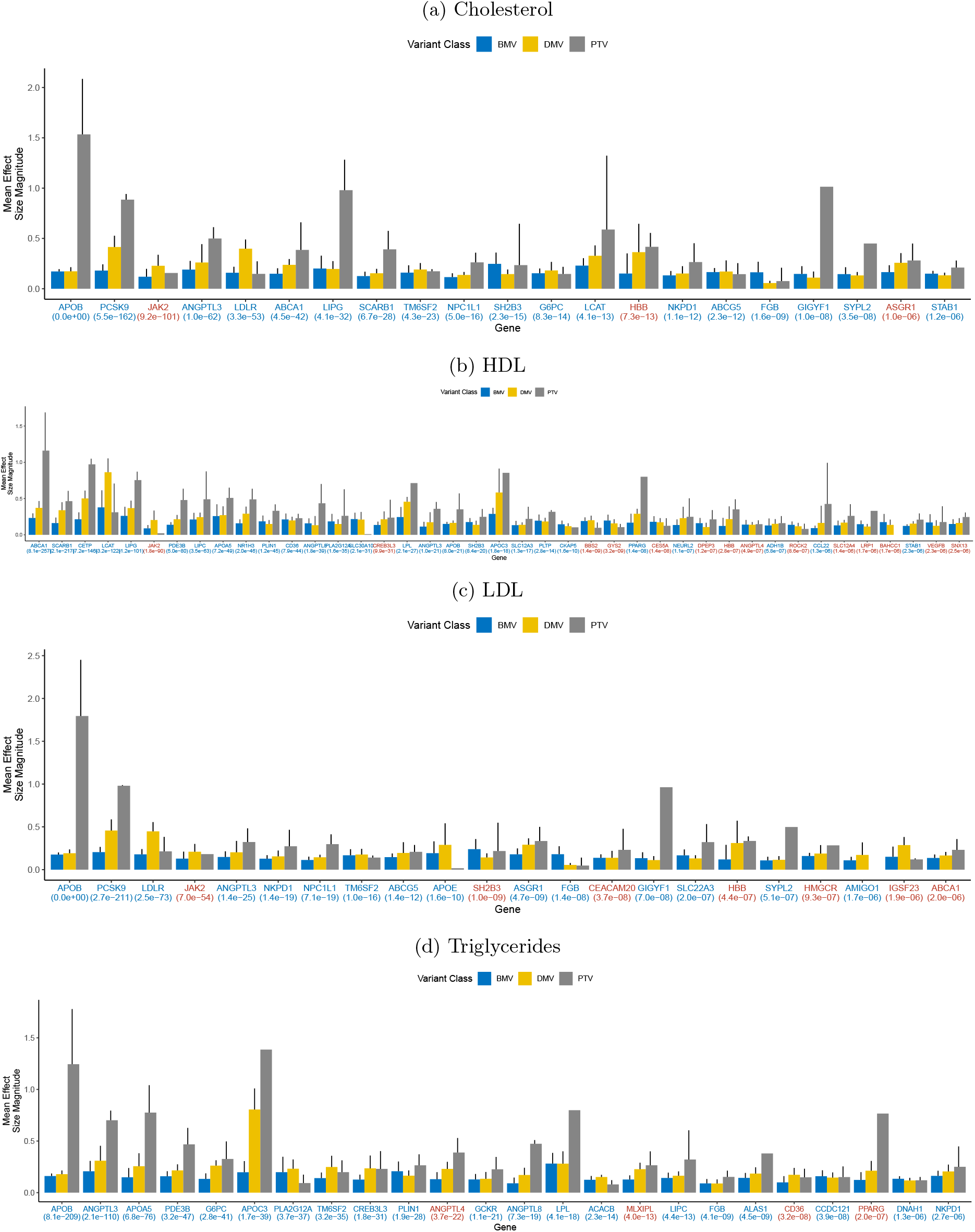
Effect size patterns among candidate lipid trait allelic series identified by COAST-SS using summary statistics from the combined MVP+UKB cohort. Shown are the mean absolute effect sizes across the BMVs, DMVs, and PTVs in a given gene. Error bars are 95% confidence intervals (CIs). The absence of a CI indicates insufficiently many variants to estimate a standard error. The gene symbol is colored blue if the gene was previously associated with the same trait via rare-variant analysis in Genebass, and red others. The number in parentheses below the gene symbol is the COAST-SS p-value.

## Discussion

In this work, we described COAST-SS, an extension of the coding-variant allelic series test that accepts commonly available GWAS-type summary statistics as input. In two respects, COAST-SS is not identical to the original COAST. First, reconstructing the allelic series max test from standard summary statistics was not possible. Rather than requiring non-standard summary statistics as input, we elected to remove the max test from COAST-SS. Second, the distinction between the count (additive) and indicator (dominance) models was dropped because investigators working from summary statistics often will not have control over how the genotypes were encoded. Nonetheless, analyses of lipid traits among the subset of 150K UKB subjects utilized for developing the original COAST demonstrate that COAST-SS is able to recover practically equivalent p-values. Moreover, extensive type I error studies based on real data (with a permuted phenotype) indicate that COAST-SS provides a valid test of association.

Although our experiments suggest that COAST-SS remains valid in the absence of LD information, we recommend providing an in-sample LD matrix whenever possible. While minimal LD is expected among rare variants, the extent to which LD can be safely ignored likely depends on accuracy with which the full sample LD matrix can be approximated by the identity matrix. In our analyses of the UKB, the LD among rare variants in the full sample LD matrix was negligible, making approximation by the identity reasonable. A limitation of the present study is that it is unclear how broadly the conclusion that LD can be safely neglected generalizes. Important directions for future research are to characterize the extent of rare variant LD in cohorts beyond the UKB, and to quantify the sensitivity of COAST-SS to unaccounted-for LD. That is, if the LD matrix is approximated by the identity out of convenience or necessity, but the true LD is non-negligible, to what extent are the operating characteristics of COAST-SS disrupted?

Contrary to expectations, utilizing pathogenicity scores informed by protein language models, including AlphaMissense, ESM1b, and PrimateAI did not appreciably improve power for allelic series detection. As illustrated in **Figure S4**, this is likely because the pathogenicity scores were generally concordant across models, and in all cases the mean pathogenicity score increased monotonically with the baseline annotation category. In an analysis of lipid traits in the UKB 390K cohort, utilizing the pathogenicity scores to adaptively set the allelic series weights provided greater power on average than reclassifying missense variants according to their pathogenicity score. A future direction is to develop models that emit allelic series weights given the collection of pathogenicity scores assigned to the rare coding variants within a gene, perhaps building on the seed-gene approach recently developed by DeepRVAT [54].

Using meta-analyzed summary statistics from the UKB and MVP, with a combined sample size of up to 840K, COAST-SS identified 108 candidate allelic series for lipid traits at Bonferroni significance. Of these, 103 (95.4%) had prior evidence supporting an association with the same trait based on rare-variant analysis in Genebass or common-variant analysis in the GWAS Catalog. Note that while the existence of an association had previously been reported, here we are reporting the potential for a dose-response relationship. Of the 5 remaining associations, all with HDL, 2 (*CREB3L3* and *HBB*) had a pattern of effect sizes consistent with an allelic series, and a review of the literature supported a role for these genes in lipid metabolism.

Our main analysis has several limitations. First, although LD information from the UKB was available for 92.3% of variants included in the meta-analysis (531K of 575K), for the remaining 7.7% of variants (44K of 575K) linkage equilibrium was assumed. To mitigate the risk of type I error inflation due to LD among the variants not represented in the UKB, the MVP summary statistics were filtered to require a MAF ≤ 0.1%, a 10-fold more stringent MAF threshold than was applied in the UKB. Second, to minimize LD differences between the populations, both the UKB and MVP summary statistics were restricted to individuals of European ancestry. Thus, lipid trait allelic series manifesting in other populations may not be represented or detected. An important direction for future research is to develop methods of testing for allelic series in the multi-ancestry setting. Third, because the summary statistics from both MVP and TOPMed have *minimum* MAFs (of ≥ 0.01% and ≥ 0.1% respectively), they have likely been selectively depleted of the largest effect size, and consequently the most informative, variants. To the extent that privacy allows, sharing summary statistics with lower MAFs would facilitate the continued development and utility of summary statistic-based methods. Fourth, although COAST/COAST-SS are tailored for detecting allelic series, they retain some power for detecting associations that do not follow a dose-response relationship. This is seen in the effect size patterns presented in **Figure 5**, although these patterns should themselves be regarded with caution because rare variant effect size estimates are based on scarce data and inherently unstable. Thus, some of the associations reported may not correspond to a genuine dose-response relationship. Fifth, the replication based on data from TOPMed, although interesting insofar as it includes summary statistics from multiple ancestries, was underpowered. Despite these limitations, the extensive overlap of our results with previously-reported lipid trait associations, and the relevance of the potentially novel HDL associations to lipid metabolism more broadly, suggest that the candidate allelic series detected by COAST-SS are promising leads.

In conclusion, COAST-SS provides researchers with a fast and easy-to-use tool for identifying rare-variant allelic series starting from commonly available summary statistics, and in doing so, will significantly expand the set of traits on which well-powered allelic series testing can be effectively performed. COAST-SS has been incorporated into the `AllelicSeries R` package, which is available on `CRAN`.

## Declaration of Interests

ZRM, RD, ST, the insitro Research Team, EF, CO, TS are employees and shareholder of insitro. JiG completed an internship at insitro.

### Acknowledgements

The authors are grateful to the participants of the UK Biobank, the MVP, and TOPMed, whose data were used with permission.

## Author Contributions

ZRM, CO, and TWS conceived of the project. TWS curated genetic and phenotypic data. ZRM, JiG, and RD developed and implemented the method. ZRM, JiG, YZ, XL, and TWS performed analyses. All authors provided scientific input. ZRM wrote the first draft of the manuscript. All authors contributed to critical revision of the final manuscript.

## Data and Code Availability

This work uses genotypes and phenotypes from the UK Biobank (https://www.ukbiobank.ac.uk/) accessed under approved application number **51766**. The coding-variant allelic series test has been implemented in the `AllelicSeries R` package, which is available on GitHub (https://github.com/insitro/AllelicSeries) and CRAN (https://CRAN.R-project.org/package=AllelicSeries).

