## Supplemental for "A Scalable Framework for Identifying Allelic Series from Summary Statistics"

January 6, 2025

### Supplemental Figures

#### Uniform QQ plots

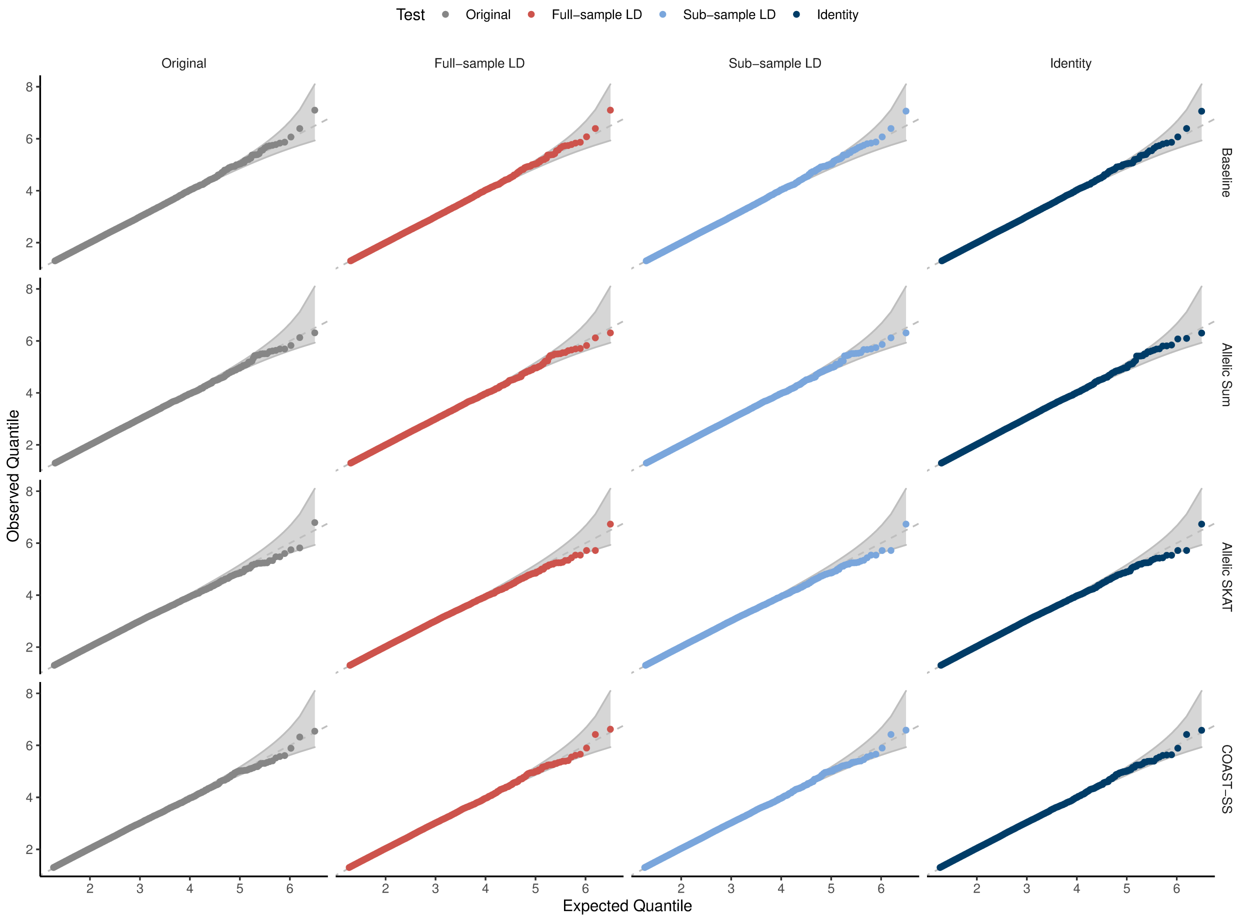


Figure S1: **Uniform quantile-quantile plots for COAST-SS and its components under various approximations to the LD matrix.** p-values were obtained using permuted LDL as the phenotype and real genotypes for 17K genes and 150K independent subjects from the UKB. The baseline, allelic series sum test, and allelic series SKAT test (top 3 rows) are composed to form the overall COAST test (bottom row). The 1st (leftmost) column provides the distribution of p-values from the original COAST, based on individual-level data, and serves as the reference. The 2nd column is COAST-SS using the full-sample LD matrix. The 3rd column is COAST-SS using a sub-sample LD matrix. The last column is COAST-SS with the LD matrix set to the identity.

### UKB 390K PP plots

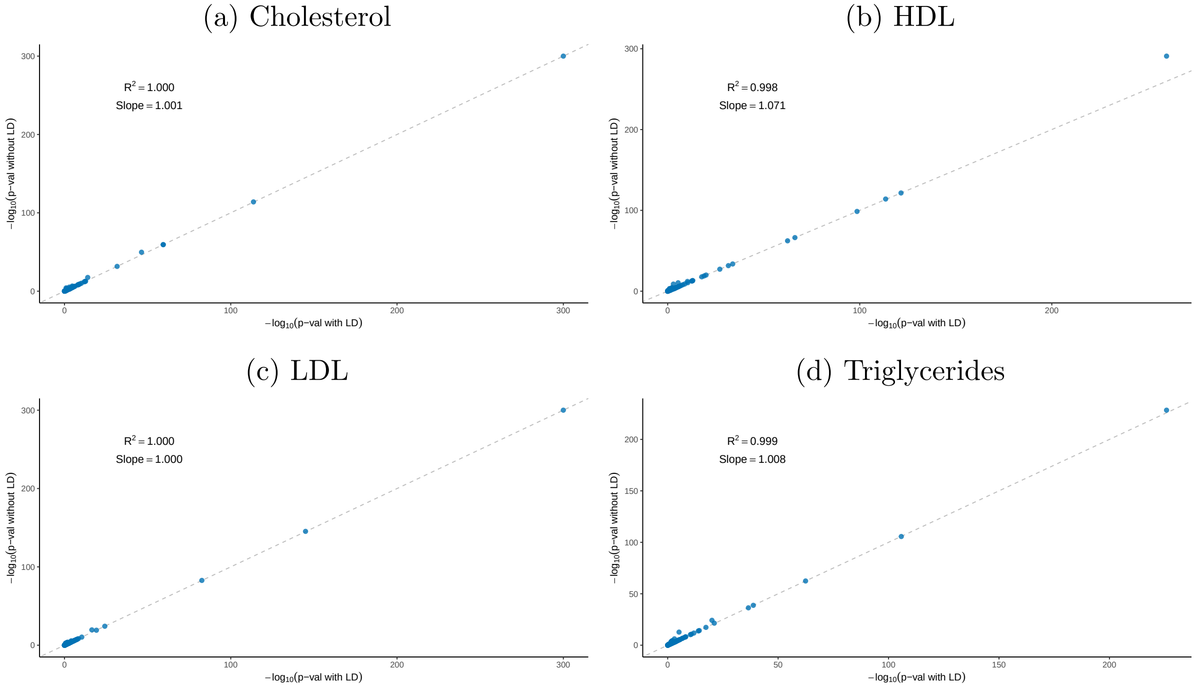


Figure S2: **Empirical concordance of COAST-SS with and without LD information as applied to lipid traits in the UKB 390K.** COAST-SS was run either with the full sample LD matrix, or with LD set to the identity. Shown are association with a $P \leq 0.05$ according to at least one of the tests.

#### SKAT power analysis

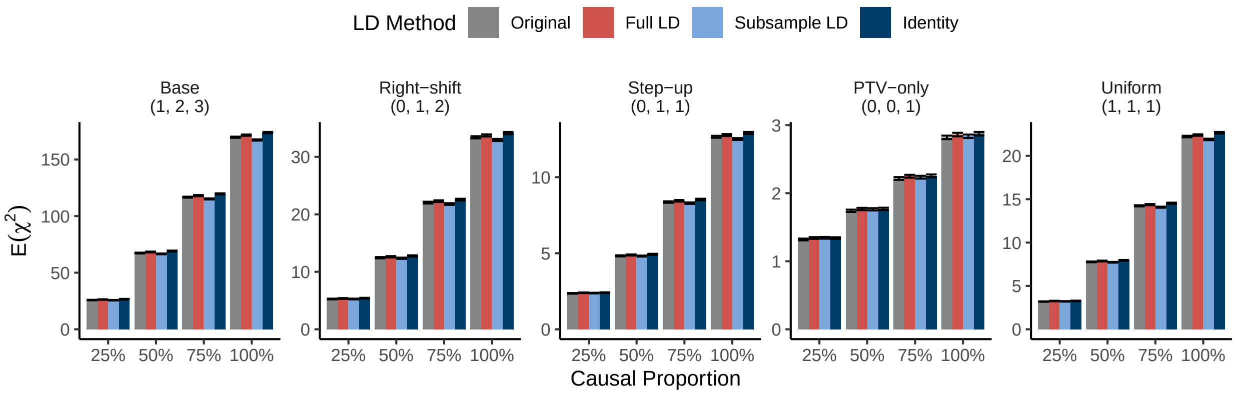


Figure S3: **COAST-SS has equivalent power to COAST, and is robust to the provided LD matrix.** Original refers to COAST run with individual-level data; full LD was calculated using all 150K subjects; sub-sampled LD was calculated using a random sample of 50K subjects, drawn with replacement; identity ignores LD. Results are presented under 5 different generative models. The tuple above each set of results indicates the true effect sizes of BMVs, DMVs, and PTVs in the generative model. Power is quantified by the expected $\chi^2$ statistic. The simulation sample size was $N = 10^4$. Error bars are 95% confidence intervals for the mean across $R = 10^5$ simulation replicates.

### Annotation and weighting schemes

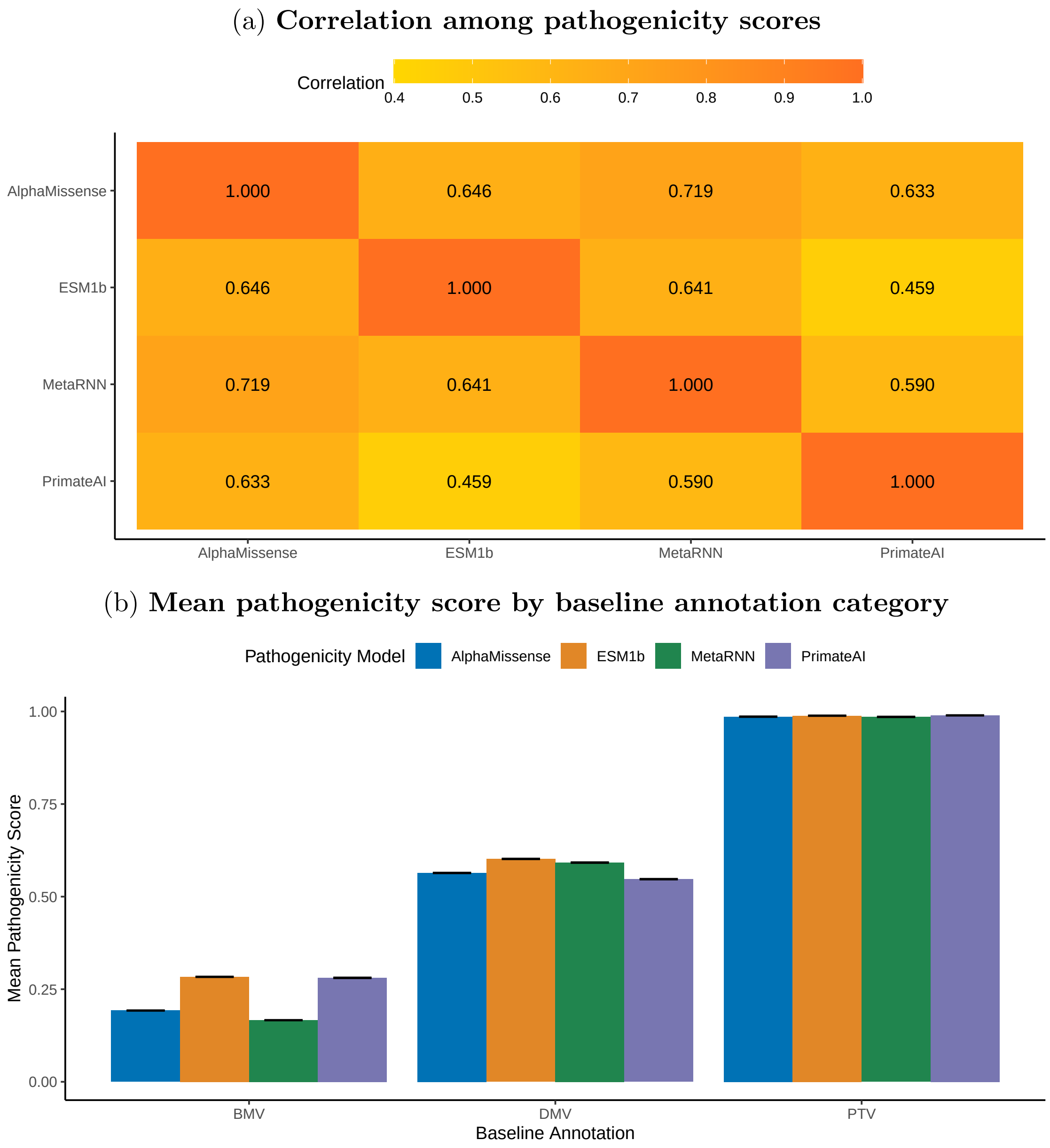


Figure S4: **Correlation among pathogenicity scores and mean score by baseline annotation categories.** (a) Correlation among the quantitative pathogenicity scores, exclding PTVs. (b) Mean pathogenicity score calculated within the baseline annotation categories BMV, DMV, and PTV. PTVs for which a pathogenicity score was unavailable were assigned the high score of 1. Error bars are 95% confidence intervals.

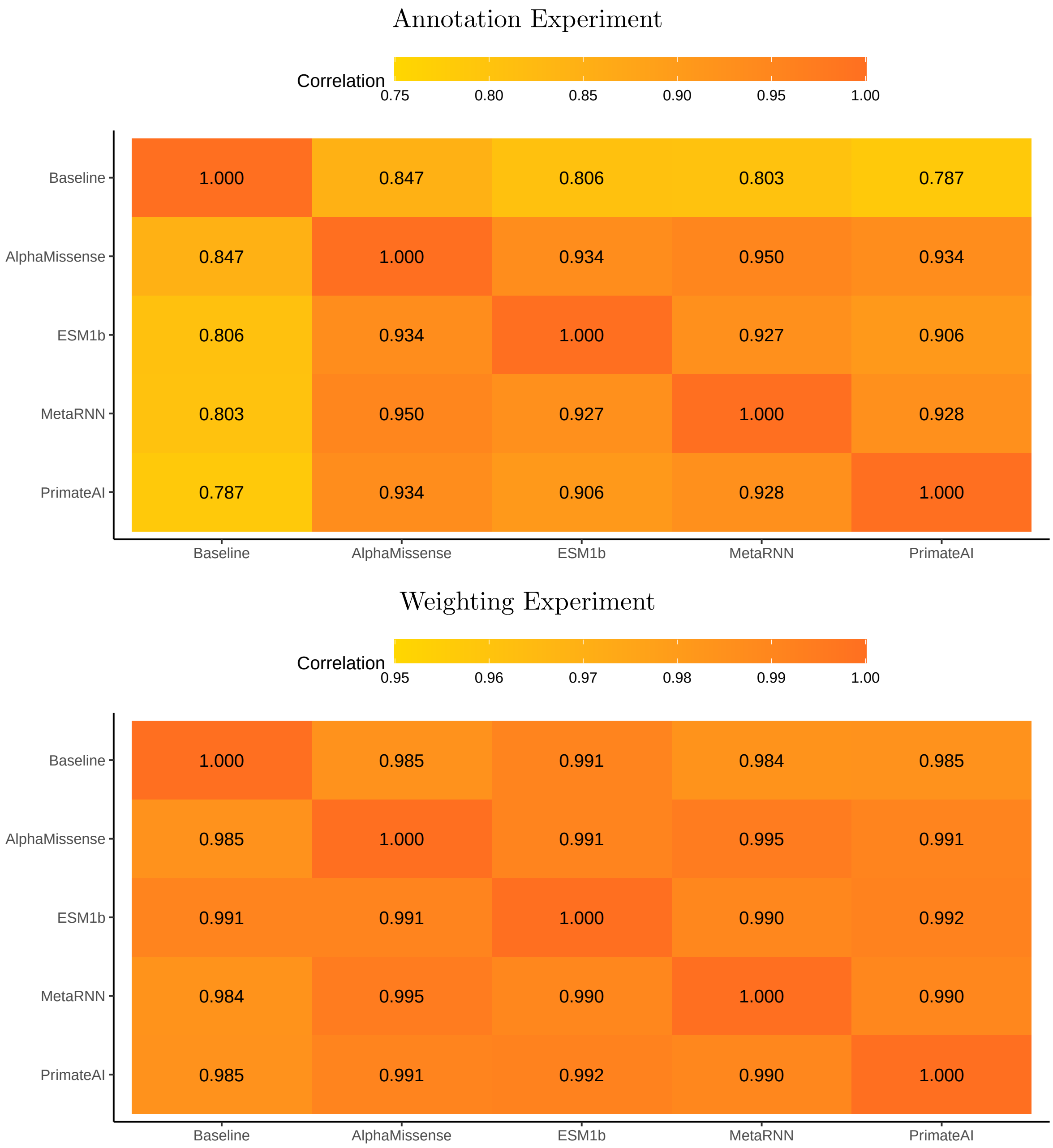


Figure S5: **P-values for lipid trait associations are highly concordant across various annotation and weighting schemes.** COAST-SS with full LD was run on summary statistics for lipid traits from 390K subjects in the UKB. Baseline is the original annotation scheme, which classifies variants into BMVs, DMVs, and PTVs. In the Annotation Experiment, missense variants were reassigned to 3 pathogenicity categories based on quantiles of a given score, while the weights were fixed at their default values, an evenly spaced increasing sequencing. PTVs were always assigned to the highest category. In the Weighting Experiment, the annotation scheme was fixed at baseline, but the allelic series weights were set, adaptively for each gene, to the mean value of the pathogenicity score for variants in that category. The value in the correlation matrix is the squared Spearman correlation between the p-values, calculated across gene-trait pairs significant according to at least 1 test.

(a) **Burden Phenotype**

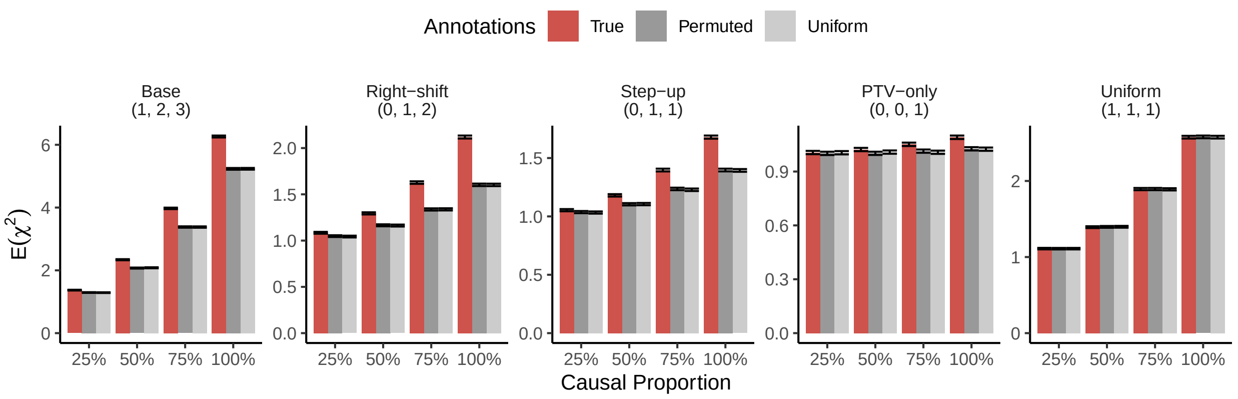


(b) **SKAT Phenotype**

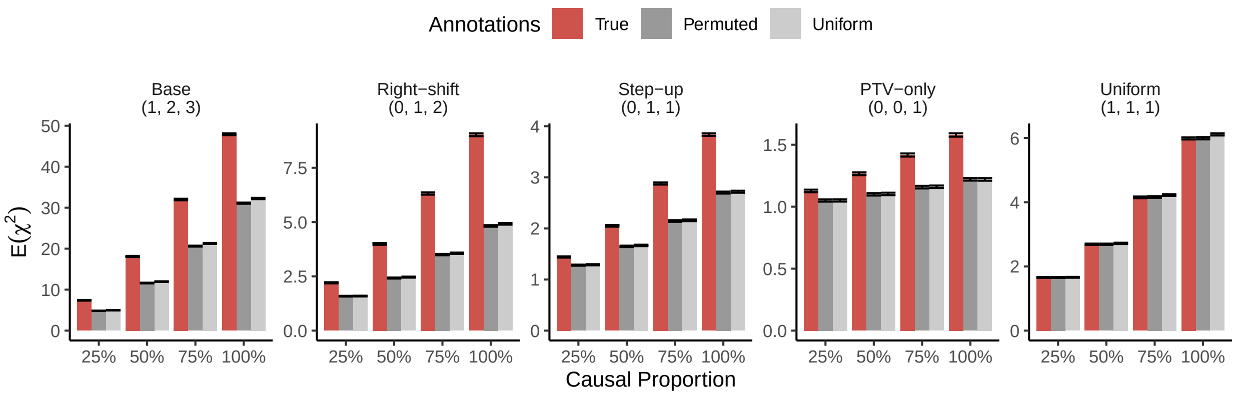


Figure S6: **Informative annotations improve power to detect allelic series.** COAST-SS was run either using the true (generative) annotation scheme, permuted annotations, or annotations sampled uniformly at random. Results are presented under 5 different generative models. The tuple above each set of results indicates the true effect sizes of BMVs, DMVs, and PTVs in the generative model. Power is quantified by the expected $\chi^2$ statistic. The simulation sample size was $N = 10^4$. Error bars are 95% confidence intervals for the mean across $R = 10^5$ simulation replicates. Note that no power gain is expected in the uniform setting (far right) because the annotations are uninformative when all categories have the same expected effect size.

#### Lipid trait analysis in UKB+MVP

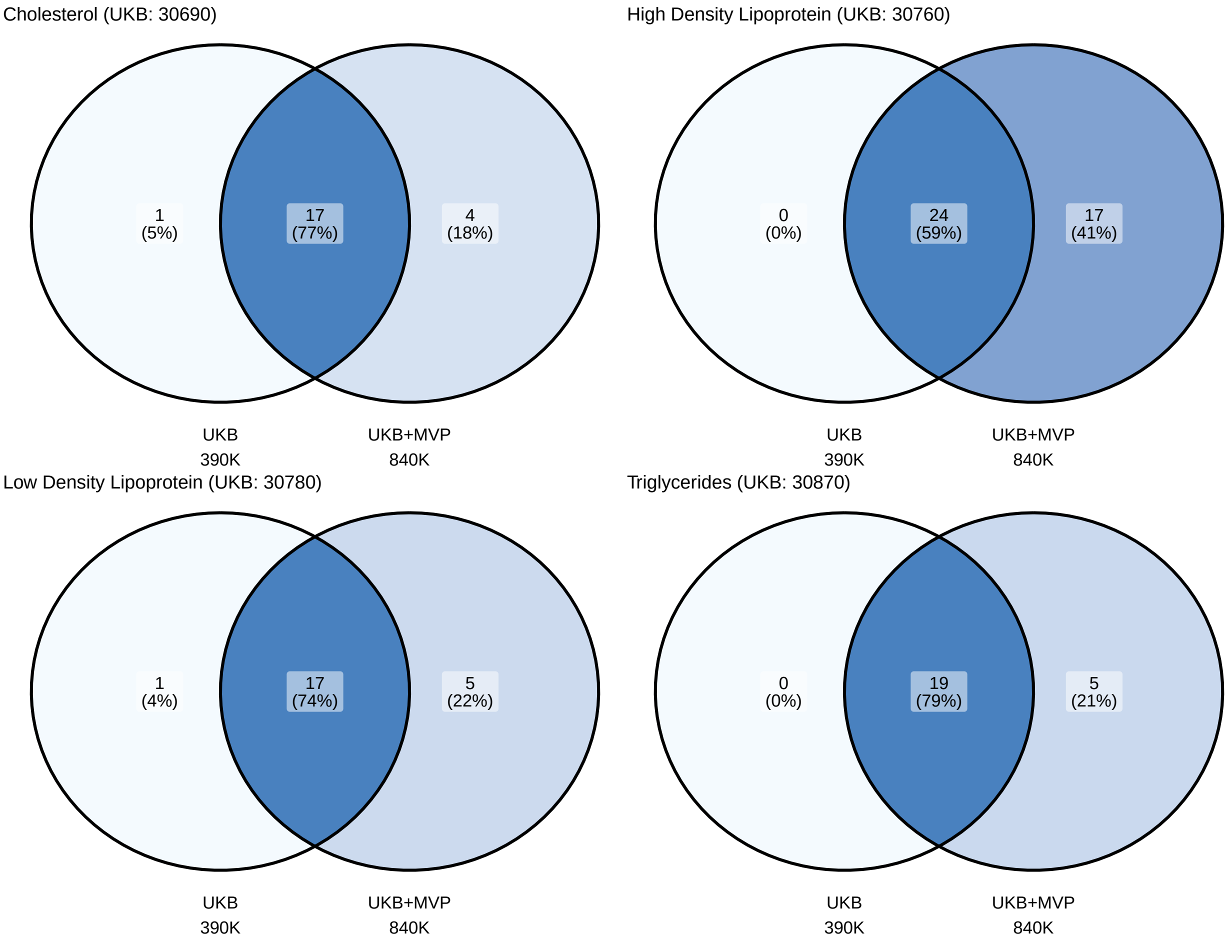


Figure S7: **Meta-analysis with MVP improves power for allelic series detection.** For each trait, left shows the number of Bonferroni significant genes detected using COAST-SS applied to summary statistics from the UKB 390K cohort, and right shows the number of Bonferroni significant genes detected in the meta-analyzed UKB+MVP cohort, with a combined sample size up to 840K.
